# The effects of both recent and long-term selection and genetic drift are readily evident in North American barley breeding populations

**DOI:** 10.1101/026625

**Authors:** Ana M. Poets, Mohsen Mohammadi, Kiran Seth, Hongyun Wang, Thomas J.Y. Kono, Zhou Fang, Gary J. Muehlbauer, Kevin P. Smith, Peter L. Morrell

**Affiliations:** Department of Agronomy and Plant Genetics, University of Minnesota, St. Paul, MN 55108; Department of Agronomy, Purdue University, West Lafayette, IN 47906; DuPont Pioneer, Mankato, MN 56001; Bayer CropScience, 407 Davis Drive, Morrisville, NC 27560

## Abstract

Barley was introduced to North America ∼400 years ago but adaptation to modern production environments is more recent. Comparisons of allele frequencies among different growth habits and inflorescence types in North America indicate significant genetic differentiation has accumulated in a relatively short evolutionary time span. Allele frequency differentiation is greatest among barley with two-row versus six-row inflorescences, and then by spring versus winter growth habit. Large changes in allele frequency among breeding programs suggest a major contribution of genetic drift and linked selection on genetic variation. Despite this, comparisons of 3,613 modern North American cultivated breeding lines that differ for row type and growth habit permit the discovery of 183 SNP outliers putatively linked to targets of selection. For example, SNPs within the *Cbf4, Ppd-H1*, and *Vrn-H1* loci which have previously been associated with agronomically-adaptive phenotypes, are identified as outliers. Analysis of extended haplotype-sharing identifies genomic regions shared within and among breeding programs, suggestive of a number of genomic regions subject to recent selection. Finally, we are able to identify recent bouts of gene flow between breeding programs that could point to the sharing of agronomically-adaptive variation. These results are supported by pedigrees and breeders understanding of germplasm sharing.

## INTRODUCTION

The evolution of breeding populations encompasses processes that reasonably mimic the evolution of natural populations, but in accelerated time (Ross-Ibarra et al., 2007). Comparative population genetic approaches can be used for the identification of genes underlying adaptive variation and to understand the effects of demographic patterns on diversity without specific phenotypic information (Nielsen et al., 2007; Ross-Ibarra et al., 2007).

Initiation and evolution of breeding populations may involve episodes such as founder events (Martin et al., 1991), bottlenecks, and gene flow from other breeding programs or exotic sources. These demographic effects can contribute to differences in allele frequency among breeding programs due to genetic drift, local adaptation, and selection (or linked selection). Understanding the selection and demographic history, including the effects of genetic drift and migration in breeding populations can accelerate crop improvement (Ross-Ibarra et al., 2007), for example by the identification of loci involved in domestication and improvement (e.g., Cavanagh et al., 2013); identification of introgression between domesticates and wild relatives (e.g., Hufford et al., 2013); and the determination of specific donor individuals contributing, for example, disease resistance variants (e.g., Fang et al., 2013).

History of selection can be investigated by the identification of large allele frequency differences among populations (at individual loci) (Cavalli-Sforza, 1966). Allele frequency differences between subdivided populations can be measured by fixation indices or *F*-statistics such as *F*_ST,_ a measure of differentiation in allele frequencies in sub-populations relative to the total population (Lewontin and Krakauer, 1973). The identification of loci with large differences in allele frequency among populations (as identified from *F*_ST_ outliers) works especially well for older or longer-term selective events (Nielsen et al., 2007).

Patterns of more recent selection (Pritchard et al., 2010; Horton et al., 2012) can be identified through approaches that detect a high degree of haplotype sharing between individuals. These approaches can identify specific haplotypes (and their underlying sequence variants) subject to selection based on their relative frequencies in a population (Hudson et al., 1994; Innan et al., 2005).

Migration between populations can contribute to adaptive variation (Slatkin, 1987). Identity by state (IBS) analysis is a sensitive approach for the identification of specific genomic regions involved in migration between distinct populations. IBS analyses have been used in studies of barley and maize to identify specific genomic regions involved in adaptation to new environments and disease resistance genes derived from exotic germplasm (Fang et al., 2013; Hufford et al., 2013).

In the present study we use single nucleotide polymorphism (SNP) genotype data from the Barley Coordinated Agricultural Project to investigate the breeding history of North American barley breeding populations. Barley (*Hordeum vulgare* ssp. *vulgare*) was introduced to North America by European colonists as early as 1602 as a crop essential for beer production (Weaver, 1950). Early successes in barley production in North America involved introduction of varieties adapted to similar environmental conditions. In the eastern growing region of the United States barley was introduced from northern Europe and England malting varieties; and in the western growing region from Mediterranean feed varieties (Weaver, 1943). Notable exceptions from outside northern Europe included “Manchuria” from northeastern China which was well adapted to the Upper Midwest growing environment, Stavropol from Russia was the most important variety in the Lower Midwest (especially Kansas), and “Trebi” from the southern shores of the Black Sea was found to be particularly productive in western regions under irrigated conditions (Wiebe and Reid, 1961). Despite the use of varieties from multiple Old World sources, the set of founder varieties was relatively narrow, which likely contributed to reduced diversity in modern North American cultivars (Martin et al., 1991).

Barley production is divided into spring and winter growth habit. Spring barley production dominates in North America, while winter barley is grown in more southerly latitudes and moderate coastal climates (Weaver, 1950). Spring and winter barley have been bred separately by breeding programs located in different geographic regions. Further trait requirements by end use markets (i.e., malting, feed, and food) and the establishment of breeding programs for two-row and six-row inflorescence type, contribute to highly structured barley populations (Cuesta-Marcos et al., 2010; Hamblin et al., 2010; Wang et al., 2012; Zhou et al., 2012).

Our analysis focuses on four major questions. First, which of the factors, including breeding programs, growth habit, and row-type contribute most directly to genetic differentiation among samples? Second, to what extent are loci identified as major contributors to phenotypic variance in Old World barley contributing to allele frequency differences in North American breeding populations? Third, can we identify evidence of recent or long-term selection acting on breeding populations and what loci are involved? Fourth, how have patterns of shared ancestry and migration contributed to diversity and relatedness in current breeding populations?

## MATERIALS AND METHODS

### Plant Materials

Genotypic data for a total of 3,971 barley accessions including varieties, advanced lines, and genetic stocks from barley breeding programs that participated in the Barley Coordinated Agricultural Project (CAP) were downloaded from The Triticeae Toolbox (T3) (http://triticeaetoolbox.org/). Barley samples are representatives of four years of germplasm enhancement (2006-2009) from 10 breeding programs. These breeding programs include most barley growing regions and market end-uses in the United States. For more information about the programs (see Hamblin et al., 2010; Wang et al., 2012).

Breeding two-row and six-row barley within a single program often involves different objectives, thus we consider these types as independent populations resulting in 16 breeding populations (see Table 1). Hereafter, the following notation to refer to each breeding population was used: “breeding program’s name abbreviation” followed by 2 or 6 for two-row and six-row, respectively as described in Table 1. The Bush Agricultural lines from the international program, referred here as BAI2, were separated from the North American two-row lines, referred here as BA2. Analyses included a total of 16 populations (Table 1). There were 10 six-row accessions from Oregon mislabeled in T3 as two-row (Dr. Patrick Hayes, personal communication), we used the corrected row-type (see Supplementary Material for more details). The current sample is divided hierarchically at the highest level into spring and winter growth habits and then within each growth habit by breeding programs (Figure 1).

**Table 1.**
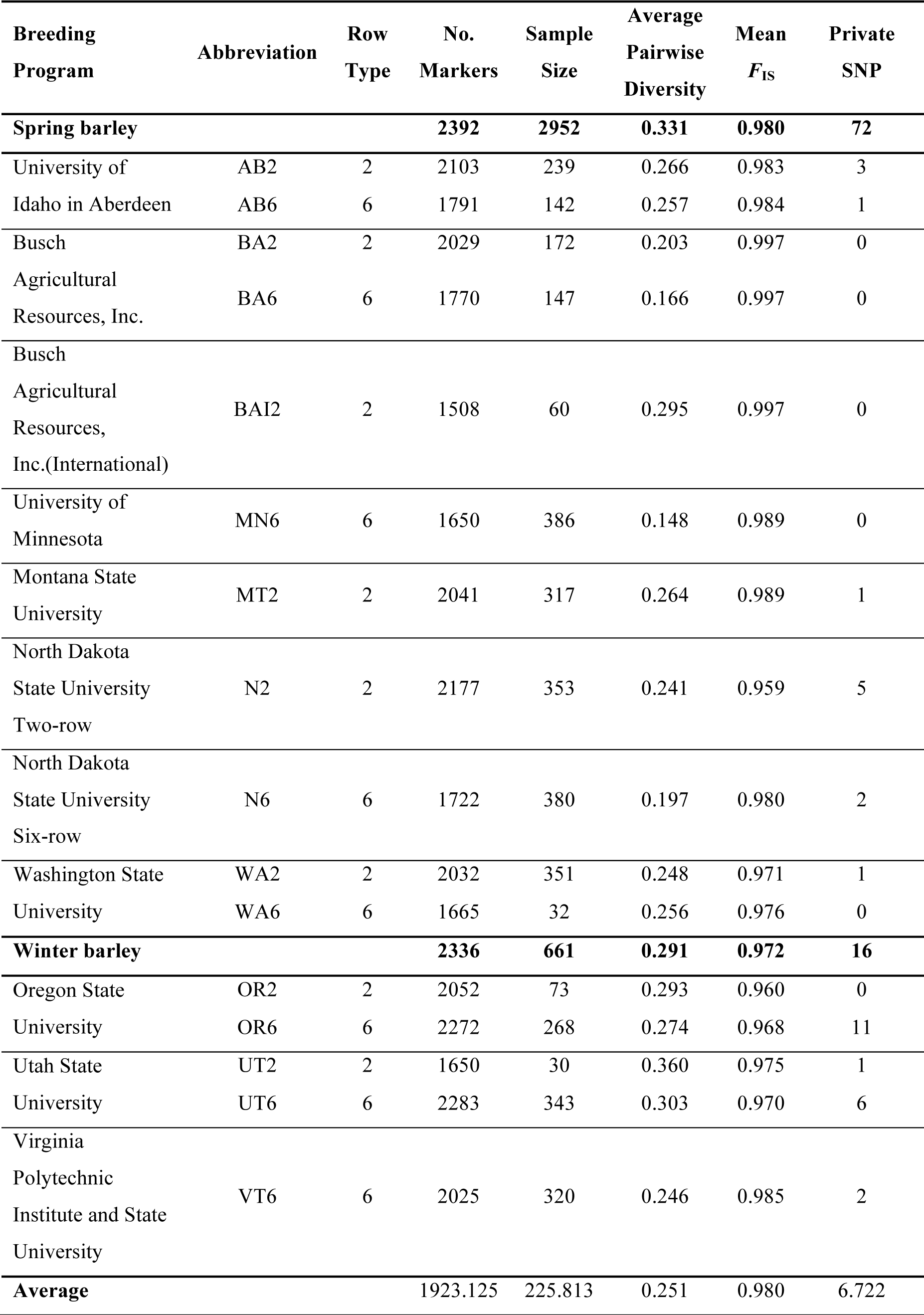
Descriptive statistics by breeding program and row type.

**Figure 1.**
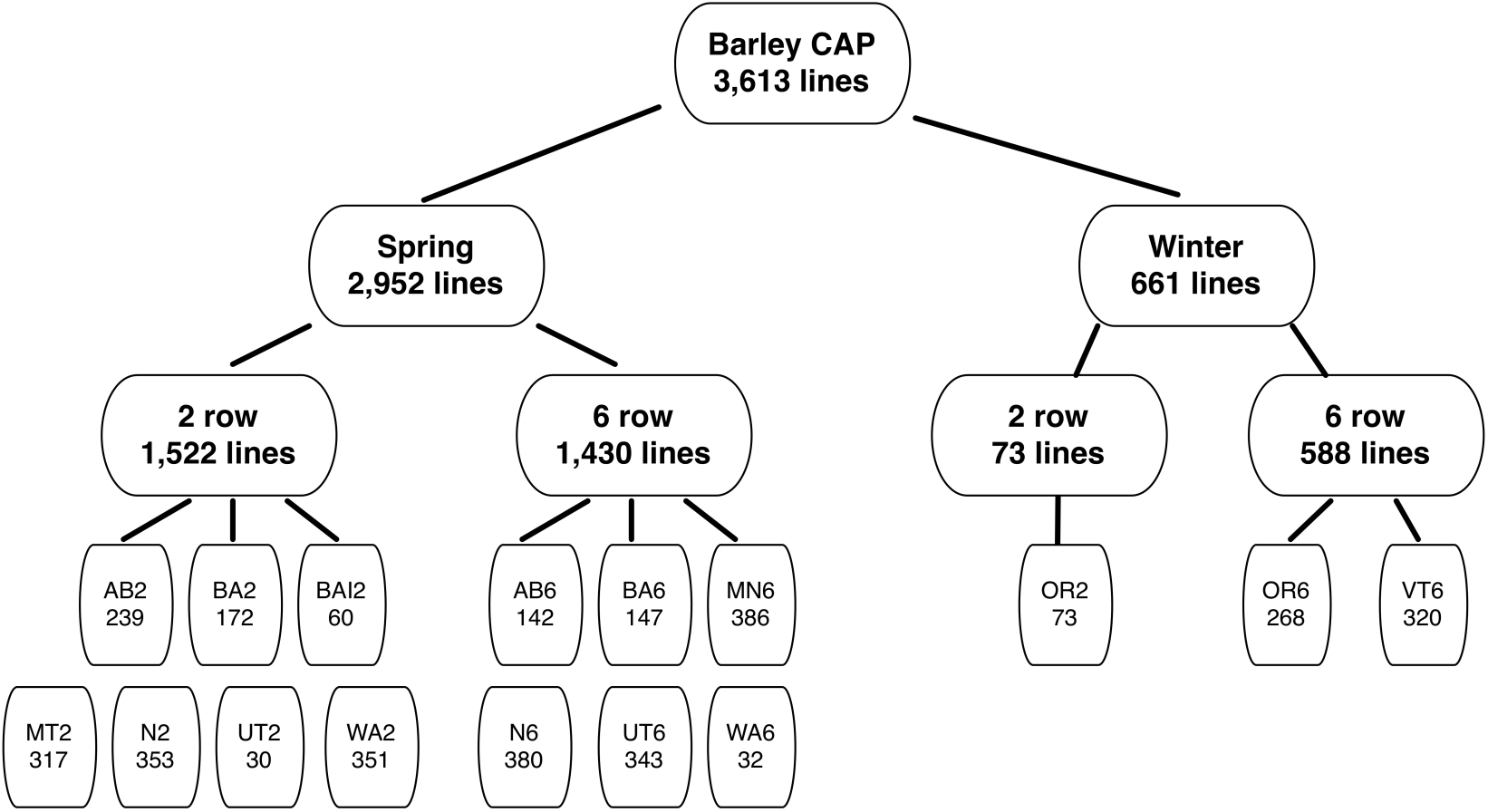
Breeding programs. University of Idaho in Aberdeen (AB), Busch Agricultural Resources, Inc. (BA), Busch Agricultural Resources, Inc. International lines (BAI), Oregon State University (OR), Utah State University (UT), Washington State University (WA), Montana State University (MT), Virginia Polytechnic Institute and State University (VT), North Dakota State University two-rows (N2), North Dakota State University six-row (N6), and University of Minnesota (MN).

### Genotyping

All 3,971 accessions were genotyped with 2,882 Barley Oligo Pooled Assay Single Nucleotide Polymorphism, referred to as BOPA SNPs (Close et al., 2009) using Illumina GoldenGate Technology (Illumina, San Diego, CA); genotypic data was downloaded from http://triticeaetoolbox.org. BOPA SNPs were identified primarily from resequencing of expressed sequenced tags (ESTs) (Close et al., 2009). SNP order along each linkage group is based on the consensus genetic map (Muñoz-Amatriaín et al., 2014). Sampled accessions were self-fertilized to at least the F_4_ generation before genotyping (Hamblin et al., 2010; Wang et al., 2012), resulting in average expected heterozygosity of 6.25%. Genotyping information was downloaded from (http://triticeaetoolbox.org/) with all filters set to zero.

SNP annotations and metadata information, including gene names and whether SNPs occur in genic or non-genic regions, were obtained using SNPMeta (Kono et al., 2013).

### Quality Control

The dataset was filtered for monomorphic SNPs, and SNPs or accessions with more than 25% missing data. Additionally, we removed accessions with incomplete sample information (i.e., row type or growth habit) and accessions where single lines represented a population. We removed accessions that presented > 6.25% heterozygosity. Finally, we removed genetic stocks or near-isogenic lines such that one accession from each near-isogenic line set was retained in the dataset.

### Summary Statistics

Basic descriptive statistics were calculated for each of the 16 breeding populations (Table 1). The degree of inbreeding was estimated by the inbreeding coefficient *F*_IS_ (1 – H_o_/H_e_) using a custom R script. The similarity between samples was estimated by the percent pairwise diversity calculated using the *compute* program from the libsequence library (Thornton, 2003). Monomorphic SNPs in each population were excluded and heterozygous and ambiguous calls were treated as missing data. The SharedPoly program from libsequence (Thornton, 2003) was used to count the number of private SNPs in each breeding population.

### Derived Site Frequency Spectrum (SFS)

To infer ancestral state, SNP states from *Hordeum bulbosum* as reported for BOPA SNPs were used (Fang et al., 2014). This involved alignment of RNAseq data from one accession of *H. bulbosum* (Cb2920/4) to the Morex draft assembly (Mayer et al., 2012), and calling the *H. bulbosum* nucleotide at the BOPA SNP positions. Ambiguous nucleotide calls, trans-specific polymorphisms, and sites at which *H. bulbosum* segregates for different nucleotides than barley (*H. vulgare ssp. vulgare)* were treated as missing data. Derived site frequency spectra were calculated and plotted for each of the 16 populations.

### Joint derived Site Frequency Spectrum

Following the same procedure as for the derived SFS for each breeding population we calculated the derived SFS within winter or spring accessions and within two-row or six-row accessions using a custom R script. The derived SFS was compared between growth habit and between row types. The joint derived SFSs were plotted using the R package grDevices (Team, 2012).

### Population Structure

The degree of differentiation among individuals from all breeding programs was estimated by Principal Component Analysis (PCA). The analysis was performed using the SmartPCA program from the EIGENSOFT package (Patterson et al., 2006). SmartPCA permits PCA analysis with SNP loci that include missing data.

### Maximum likelihood tree of relatedness and migration

The population relatedness and patterns of gene flow between breeding populations was inferred using a maximum likelihood approach implemented in TreeMix (Pickrell and Pritchard, 2012). To polarize the divergence among populations we used genotyping data for 438 landrace accessions from Europe (Poets et al., 2015) (see https://github.com/AnaPoets/BarleyLandraces). The majority of founders of the North American barley population derive from Europe (Weaver, 1944). The 2,021 SNPs shared between data sets were used to build a phylogeny tree using landraces as an outgroup. We ran 25 replicates of the tree, bootstrapping with 75 SNP windows. We used the replicate with the lowest standard error for the residuals as the base tree topology and inferred the likelihood of having between one and five migration events among breeding programs. The plot of the residuals was used to evaluate which tree best fit the data. Candidates for admixture can be identified by those population pairs with residuals above zero standard error, which represent populations that are more closely related to each other in the data than in the best-fit tree (Pickrell and Pritchard, 2012).

### Changes in allele frequency

To identify putative targets of long-term selection involved in the row-type and growth habit differentiation among breeding programs we used the Weir and Cockerham (Weir and Cockerham, 1984) measure of allele frequency differentiation, *F*_ST_, as implemented in the R package hierfstat (Goudet, 2005). *F*_ST_ was calculated for the following partitions of the data: 1) spring versus winter, 2) two-row versus six-row, and 3) among breeding programs. Owing to the high level of inbreeding in the data set, a haploid model for *F*_ST_ estimation was used. Heterozygous SNPs were treated as missing data. An empirical genome-wide threshold for the top 2.5% of *F*_ST_ values was used to identify SNPs with large differences in frequency relative to the genome-wide average. To identify the degree of differentiation that each breeding program has with respect to other programs we report *F*_ST_ for all pairwise comparisons.

To characterize average allele frequency divergence for breeding populations an analogous to *F*_ST_ was calculated according to Nicholson *et al*.,(Nicholson et al., 2002) and reported as *c*. The *c* is an estimate of the degree of divergence of a population from ancestral allele frequencies. For this analysis, the data set was divided into two groups: spring six-row, and spring two-row. We do not report *c* for winter barleys owing to limited sampling. Monomorphic SNPs were removed from each group. *c* was calculated using the popdiv program from the popgen package in R (Marchini, 2013). We used a burn-in period of 1,000 iterations followed by a run length of 10,000 iterations with the scale parameter of Dirichlet distribution used to update global allele frequencies *m* = 10 (see Supplementary Material for more details).

### Analysis of resequencing data for known genes contributing to phenotypic differentiation

Resequencing data for 10 accessions for vernalization sensitivity loci (*Vrn-H3*) (von Zitzewitz et al., 2005) and 96 accessions for *Vrs1* gene controlling row-type differentiation (Komatsuda et al., 2007) were obtained from NCBI Popsets (UID #157652625 and 219664771). Contextual sequences for SNPs known to occur in these genes were downloaded from T3. To determine the position of SNPs within these genes and their correlation with growth habit and row-type differentiation SNP contextual sequence for individual SNPs were aligned to resequencing data set in Geneious v.7.1.9 (Kearse et al., 2012).

### Identity by State

We used an identity by state (IBS) analysis to identify shared genomic segments between breeding programs potentially indicative of recent introgression. The analysis used PLINK v.1.90 (Chang et al., 2015) with windows sized of 50 and 100 SNPs allowing for up to 10% mismatch. The frequency of shared segments between two populations was estimated for each SNP window. This analysis made use of phased genotyping data, with phase inferred using fastPHASE v1.2 (Scheet and Stephens, 2006). Missing genotypic state was treated as missing (*i.e.,* genotypic state inferred during phasing were ignored). The phased data were only used for IBS and pairwise haplotype sharing analyses.

### Pairwise Haplotype Sharing

To explore recent events of selection within populations, we used the pairwise haplotype sharing (PHS) approach (Toomajian et al., 2006). A shared haplotype is defined as a genomic segment that extends out from a focal SNP, and is shared among individuals in a population. PHS is a form of IBS analysis that compares the extent of shared haplotypes among individuals normalized by genome-wide sharing. A PHS score depends on the length of the shared haplotype and its frequency in the population. Extended shared haplotypes are potentially suggestive of recent or ongoing selection, owing to limited potential for recombination to break down genomic regions subject to recent selection (Horton et al., 2012). PHS was calculated within each breeding population using a customized Perl script (Cavanagh et al., 2013). An empirical threshold of 2.5% and a minimum allele frequency of 10% within each population were used to identify outliers in the distribution of PHS.

### Four-population test for gene flow detection

The robustness of patterns of migration inferred using TreeMix (see Results for details) was assessed using the four-population test *(f_4_-test)* (Keinan et al., 2007; Reich et al., 2009). The *f*_*4*_-*test* is designed to distinguish introgression from incomplete lineage sorting. The test evaluates trees of relatedness among populations and measures genetic drift along lineages quantitatively based on the variance in allele frequencies. Significant deviations from zero in three possible tree topologies (see Supplementary Material for details) indicate that the tree evaluated does not fit the data, suggesting the presence of gene flow.

We used the topology inferred in TreeMix as our hypothesized relationship between populations. We inferred that a tree of relatedness with three migration events fit the data better than having more or less migrations (see the Results section for more details). Following the ((A, B),(C,D)) notation for a tree topology with four populations (Figure S1), we assessed migration among the following sets of populations: ((N2, X),(Y, Y)), ((UT6, UT2), (Y, Y)) and ((OR2, X), (OR6,VT6)), where X and Y were replaced iteratively for any two-row or six-row barley populations, respectively. The populations UT6, OR2 and N2 were chosen because they had more specific signal of gene flow according to the TreeMix results. These populations were paired with populations from the same branch of the tree inferred by TreeMix, assuming that these populations are more similar to each other due to shared ancestral polymorphisms. The other pair of populations to be compared to was selected from the branch containing the population putatively involved in the migration event (connected by the arrow, Figure 6). Since the population putatively involved in migration was not clearly defined, we ran the analysis iteratively between pairs of populations taken from the same branch. We used the fourpop option in the TreeMix software (Pickrell and Pritchard, 2012) to estimate the *f*_*4*_-*value* for each configuration. Significance of *f*_*4*_-*values* was determined at p <0.05. We infer that migration has occurred when the three possible trees had a significant non-zero *f*_*4*_-*value.*

All code used for analysis and figures is available at https://github.com/MorrellLAB/NorthAmerica_Fst

## RESULTS

The original dataset included 3,971 accessions genotyped with 2,882 SNPs representative of 10 breeding programs across the United States of America. We removed 340 SNPs monomorphic in the full panel, 241 accessions with ≥25% missing data, 22 accessions with heterozygosity > 6.25% across SNPs, 13 accessions with missing growth habit (spring versus winter) or row type (two versus six rows) information, 79 near isogenic lines, and three accessions with single accessions representing a population. After quality control our data set consisted of 3,613 barley accessions (Table S1) and 2,542 SNPs (Table S2).

### Characterization of North American breeding programs

Analysis of the structure of the North American populations using principal component analysis (Figure S2) revealed that the primary population structure (PC1 = 19.6% variance) is explained by differences in row-type, which corresponds to an average *F*_ST_ of 0.23 (Figure S3). PC2 indicates that 9.1% of the variance among lines is explained by differentiation in growth habit, with average *F*_ST_ of 0.17. These results are congruent with earlier analyses on a subset of these populations (Cuesta-Marcos et al., 2010; Hamblin et al., 2010; Wang et al., 2012) and on a comparable set of samples analyzed by Zhou *et al.* (Zhou et al., 2012).

The 16 breeding populations (*i.e.,* breeding programs separated by row-type) were represented by an average sample size of 225 lines with a minimum of 30 lines (from UT2), and a maximum sample size of 386 for MN6 (Table 1). On average, only two SNPs were private to each of the breeding programs, with a maximum of 11 private SNPs in OR6. Winter programs had 16 private SNPs respect to spring programs which in turn had 72 private SNPs (Table S3). The average inbreeding coefficient (*F*_IS_) across populations was 0.98 as expected after four generations of self-fertilization. Percent pairwise diversity ranged from 0.15 to 0.36 and averaged 0.25 across breeding programs (Table 1, Figure 2A).

**Figure 2.**
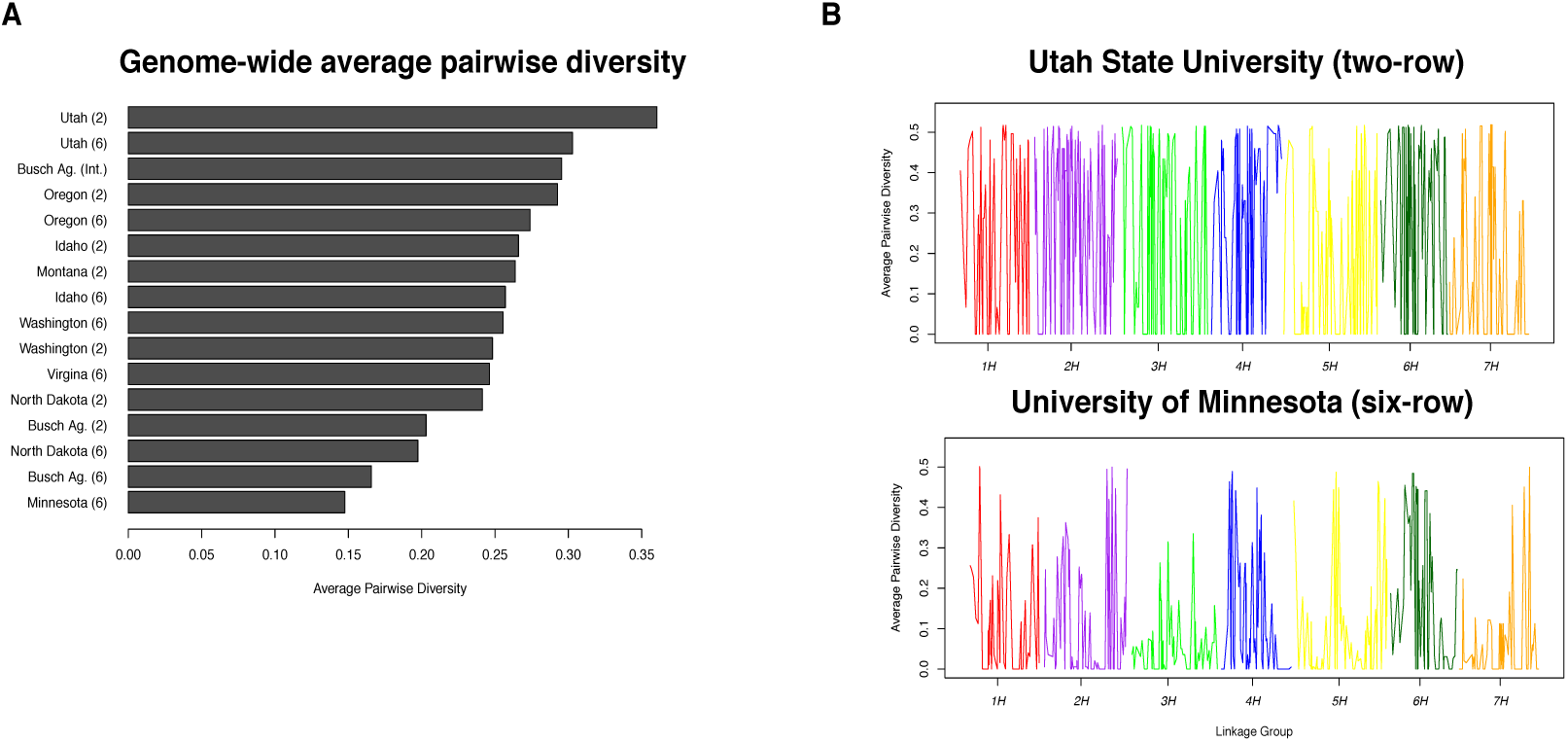
Genetic diversity in breeding programs. (**A)** Average pairwise diversity in each breeding program. (**B)** Genome-wide pairwise diversity for the most and least diverse populations according to average diversity, Utah State University (two-row) and University of Minnesota (six-row), respectively. Diversity values were averaged in 10-SNP sliding windows with a step of five SNPs.

The joint unfolded site frequency spectrum showed that there are slightly more rare variants in spring than winter programs and slightly more rare segregating variants in six-row than in two-row programs (Figure S4), reflecting minimal impact of ascertainment bias in these partitions. In individual breeding populations there is an excess of rare and high frequency variants relative to neutral expectations on a model of a population in equilibrium (Kimura, 1983) (Figure S5). OR2, OR6, UT6 and UT2 have a higher proportion of mid-frequency variants than other populations. When the breeding populations are considered jointly, the derived SFS (Figure S6) displays an elevated number of mid-frequency variants, consistent with retention of variants segregating at an average minor allele frequency of 24% in the discovery panel (Close et al., 2009).

### Genome-wide scan for evidence of selection in the North America breeding programs

The distribution of *F*_ST_ statistics for comparisons of growth habit, row-type, and breeding programs showed that among the three classifications, the greatest differentiation in allele frequencies is found among breeding programs with an average *F*_ST_ of 0.37, while 0.23 for inflorescence type and 0.17 for growth habit (Figure 3, Figure S3). For each of the three partitions, we identified 61 SNPs in the upper ≥0.975 of *F*_ST_ values, for a total of 183 SNP outliers (Table S4, S5, S6), 34 SNPs were common outliers between the row-type and breeding program comparisons, and seven were common between breeding program and growth habit comparisons.

**Figure 3.**
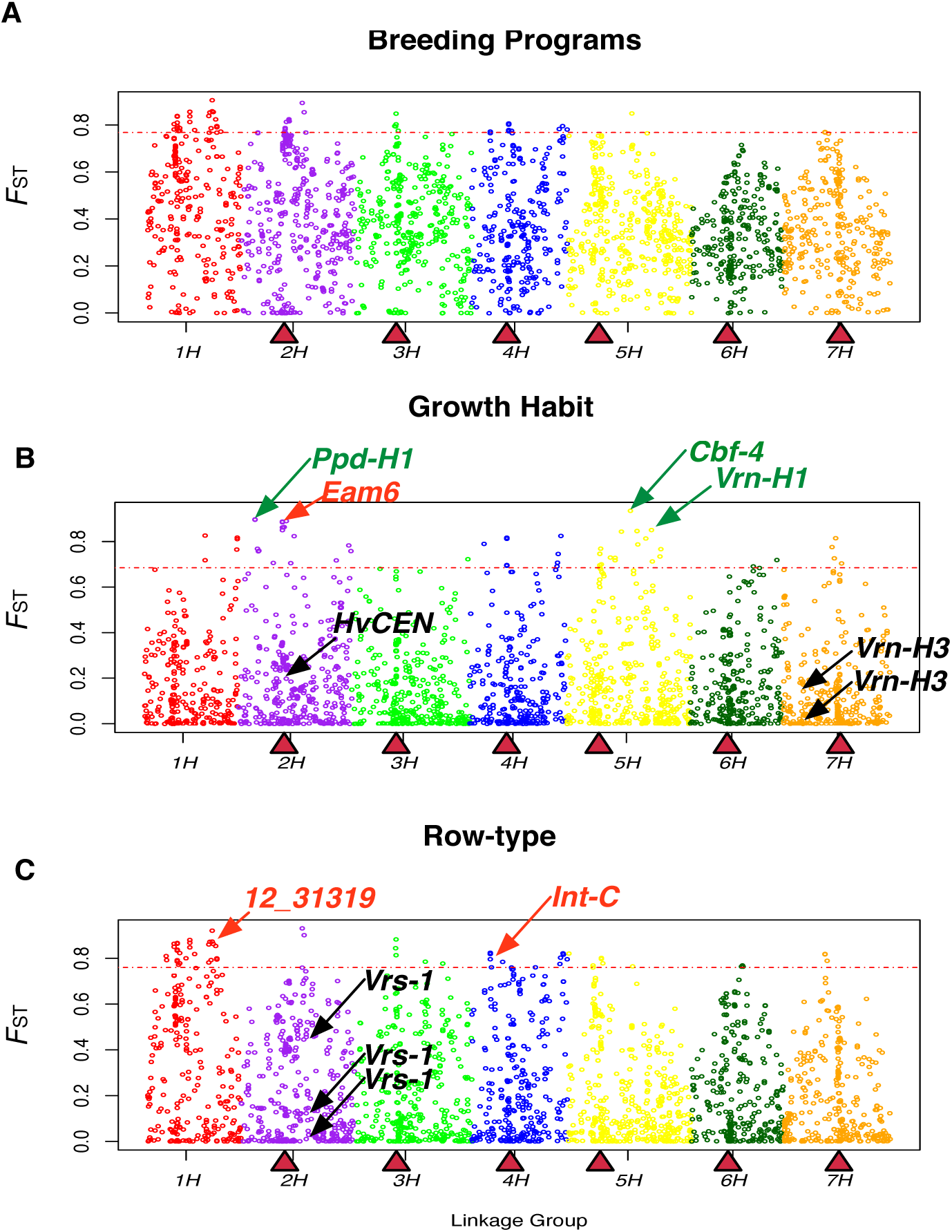
Distribution of *F*_ST_ values from comparisons between different partitions of the data. **(A)** Between spring and winter types; and **(B)** between six-row and two-row types. Triangles indicate the genetic position of centromeres (Muñoz-Amatriaín et al., 2011). Dotted line corresponds to the 97.5 percentile of the *F*_ST_ values distribution. Linkage groups are shown with different colors. In green font are genes corresponding to outlier *F*_ST_ SNPs. In red font are either the SNP linked to a predicted gene involved in the phenotype or the gene itself. The arrows below the threshold line indicate SNPs located in genes known to be involved in the phenotype.

### Known genes contributing to phenotypic differentiation

We identified a SNP outlier (12_30883, linkage group 5H) in the *F*_ST_ comparison for growth habit located in the vernalization sensitivity locus (*Vrn-H1*) (von Zitzewitz et al., 2005) known to be involved in the growth habit differentiation. SNPs within two additional well characterized genes photoperiod response-H1 (*Ppd-H1*) (Turner et al., 2005; Jones et al., 2008) and c-repeat binding factor 4 (*Cbf-4*) (Haake et al., 2002) were also found to be *F*_ST_ outliers (Figure 3B). Both of these genes have reported functions related to growth habit differentiation. *Ppd-H1* alters flowering time, thus making it possible to avoid extreme unfavorable seasonal conditions (Lister et al., 2009) whereas *Cbf-4* contributes to cold acclimation (Skinner et al., 2005).

While vernalization sensitivity and photoperiod response are important determinants that contribute to growth habit through an environmental response, there are also loci that contribute to flowering time independent of environmental cues. This class of genes in barley is referred to as Earliness *per se* (EPS) loci; one characterized example is the early maturity 6 locus (*Eam6)* (Boyd et al., 2003). A linkage mapping study placed early maturity 6 locus (*Eam6*) (Laurie et al., 1995) near the centromere of linkage group 2H (cM 67.8), a region where we identify six SNPs with elevated *F*_ST_ (average *F*_ST_= 0.87) (see Table S5, Figure 3). The two SNPs (11_20438 and 11_20366) most strongly associated with flowering time in a recent GWAS study (Comadran et al., 2011) that helped to the identification of the barley *Centroradialis* gene (HvCEN) responsible for flowering time variation (Comadran et al., 2012) were not identified as outliers in our *F*_ST_ comparison of growth habit. However, both SNPs, 11_20438 and 11_20366, have an above average *F*_ST_ 0.52 and 0.25, respectively.

In an association mapping (AM) study Cuesta-Marcos *et al.* (2010) identified two significant SNPs in linkage groups 1H and 2H (12_31319 and 11_10213, respectively) associated with row-type differentiation. In our analysis, 12_31319 has an outlier *F*_ST_ value of 0.86, while 11_10213 has an above average *F*_ST_ = 0.47 but is not defined as an outlier (Figure 3C). Additionally, there are two SNPs (11_20422 and 11_20606) in linkage group 4H that have been identified near the *Intermedium-C* gene (*Int-c*) (Ramsay et al., 2011), which is a modifier of lateral spikelets in barley. After quality control, only 11_20422 remained in our dataset. This SNP has a *F*_ST_ value of 0.81 and is thus considered an outlier in the comparisons between row types (Figure 3C).

Other SNPs occurring in well-characterized genes did not appear as outliers despite the previously reported contribution to function. There were three SNPs in our data set (12_30893, 12_30894, and 12_30895 on linkage group 7H) occurring within the vernalization sensitive locus 3 (*Vrn-H3)* (Yan et al., 2006) that were not outliers in the *F*_ST_ comparison between spring and winter types (Figure 3). This gene is an ortholog of the *Arabidopsis thaliana* flowering locus T (*FT*) that promotes flowering time under long days (Turck et al., 2008). In barley, nine linked polymorphisms in the first intron have been predicted to be responsible for the variation in flowering time at this locus (Yan et al., 2006). Alignment of the three outlier SNPs to resequencing data of this gene from 10 barley accessions (Karsai et al., 2008) identified three of the SNPs segregating in the first intron of *Vrn-H3* (Figure S7A). In Yan *et al.* (2006) nucleotide states “A” and “G” in 12_30894 and 12_30895, respectively, were associated with spring barley types, while, “T” and “C” with winter types. The resequencing data in part support this association having two out of four spring barleys with the inferred haplotype while five out of six winter barleys carried the correct inferred haplotype. However, in our larger data set of spring and winter accessions these SNPs are segregating at an average allele frequency of 50% in both spring and winter growth habits (Figure S7B), showing no association between these SNPs and spring versus winter growth habit (maximum *F*_ST_ = 0.04).

The *Vrs1* gene, a well characterized contributor to row-type (Komatsuda et al., 2007) was not identified in the *F*_ST_ comparison between two-row and six-row accessions. Our SNP panel included five SNPs (12_30896, 12_30897, 12_30899, 12_30900, and 12_30901, linkage group 2H) in *Vrs1*. However, none of the SNPs reach the empirical cutoff for *F*_ST_ in this comparison, with a maximum observed *F*_ST_ of 0.45. The “G” nucleotide state at SNP 12_30900, results in an amino acid substitution associated with the six-row phenotype (*vrs1.a3* allele), however, the alternative nucleotide state “C” can be found in either six-row or two-row accessions (Komatsuda et al., 2007; Youssef et al., 2012). In a panel of 96 European accessions of cultivated barley (Popset ID 219664771) (Figure S7 B) the “G” state always resulted in a six-row phenotype. In our sample of North American breeding programs 2% of individuals that carry this variant state were reported as two-row barleys, which is similar to previous results that found this SNP significant for row-type differentiation segregating in two-row accessions with an allele frequency of 1% (Cuesta-Marcos et al., 2010). The “C” state segregated in 50% frequency in each of the row-type partitions.

### Haplotype sharing and evidence for recent selection

The pairwise haplotype sharing (PHS) analysis permits the identification of genomic regions that are putatively involved in more recent selection. PHS analysis within individual breeding populations identified a total of 775 SNPs in the upper ≥0.975 of the PHS distribution (Table S7, FigureS8). In a small number of cases, focal SNPs in the PHS analysis were identified as outliers in more than one breeding population. Sharing of PHS outliers is greatest for OR6 and UT6 (22), BA6 and WA6 (17), and AB2 and OR2 (14) (Table S8). Average values were considerably lower, with three SNP shared within two-row populations, four within six-row populations, and two SNPs between two-row and six-row populations. Out of the 775 SNPs, 77 were in genes with known function. The haplotypes for these significant PHS values varied in length from 19.7 cM to 139.6 cM with mean 55.9 cM across breeding programs (Figure S9A, Table S9). The frequency of the SNP state with significant PHS ranged from 10% (the minimum value we consider) to 61% with mean 24% (Figure S9B).

Within BAI2, BA6, MN6, UT2, UT6, WA6, and OR2 we observed long runs of haplotype sharing (average length 112.45 cM) at an average frequency of 24%, significantly exceeding genome-wide similarities. These regions were putatively subject to recent selection. Among outliers for PHS, BA2 showed significant PHS surrounding three SNPs (12_30893, 12_30894, 12_30895, in linkage group 7H) with all three SNPs occurring within *Vrn-H3*. AB2, MT2, OR2 and OR6 had an outlier PHS value for SNPs 12_30901 (linkage group 2H) in the *Vrs1* gene. The SNPs in *Vrn-H3* and *Vrs1* were at an average frequency of 22.25 % in each of these populations, with an average length of shared haplotype of 66.98 cM (Table S7 and S9). There are three SNP (12_20368, 12_20593, and 12_21049 in linkage group 2H) with significant PHS value in OR2 (frequency 11 % and haplotype length 138.9 cM). SNPMeta annotations identified SNP 12_20593 within the nicotianamine synthase 2 (*nashor2*) gene in barley (Herbik et al., 1999) and SNP 12_20368 within a gene with sequence similarity to galactinol synthase 2 gene in wheat (*TaGolS* 2), which in turn is orthologous to the characterized gene *TaGolS* 2 in *Arabidopsis thaliana* (Taji et al., 2002). In *A. thaliana, TaGolS* 2 has been identified as playing an important role in drought-stress tolerance (Taji et al., 2002).

### Drift from an ancestral allele frequency

Quantification of allele frequency divergence based on *c* (Nicholson et al., 2002) showed that six-row breeding programs have experienced more divergence than two-row programs, with mean *c* of 23.1% and 17.1%, respectively (Figure 4, Table S10). Among the two-row programs Utah two-row has diverged the most while Idaho two-row resembles ancestral allele frequencies, suggesting reduced effects of drift or linked selection in the latter population. Among the six-row programs Utah six-row was the closest to ancestral allele frequencies while Minnesota six-row has experienced the most divergence.

**Figure 4.**
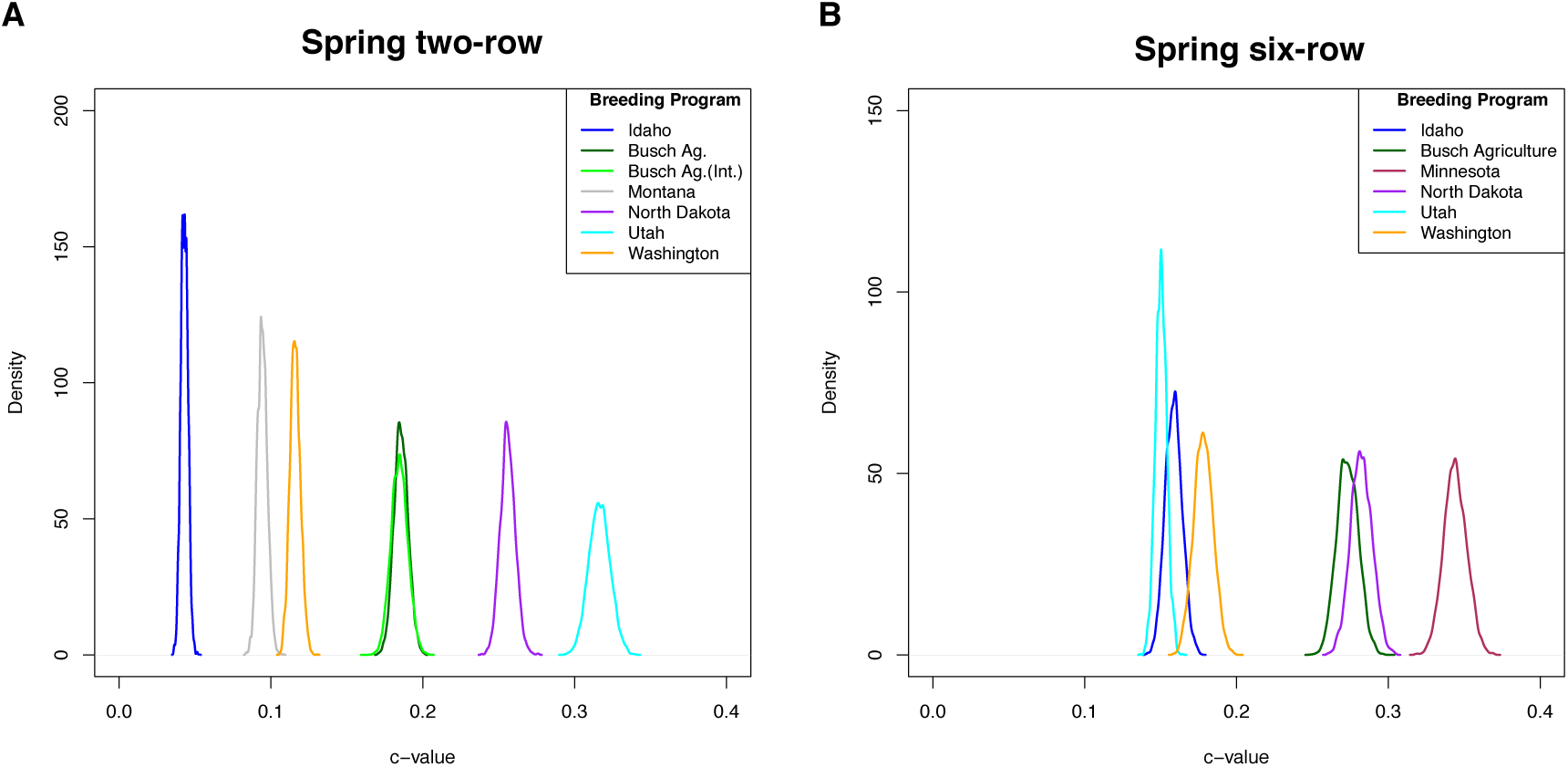
Marginal posterior density plots obtained for c for spring barley populations. Density distribution after 10, 000 iterations. Plot shows the amount of drift estimated from ancestral allele frequencies (0,0 coordinates). **(A)** Two-row barley populations; **(B)** Six-row barley populations.

### Gene flow between breeding programs

To identify genomic segments subject to recent introgression, we tested for regions with a high degree of identity by state between breeding populations. Window sizes of 50 (0.63 - 44.96 cM) and 100 (13 – 62.25 cM) SNPs were used, and we permitted up to 10% mismatch among haplotypes. As expected, populations within growth habit and row-type had a higher degree of haplotype sharing than between these partitions. The degree of haplotype sharing was lower within row-type than within growth habit (Table S11 and S12, and Figures S10 and S11). There was a high frequency of shared haplotypes among two-row populations at 50 SNPs windows, but this was reduced when 100 SNPs windows were considered. At 100 SNPs windows BAI2 and N2 presented the lowest degree of shared haplotypes with other two-row populations, while BA2 shared the most haplotypes (Figure S11C,E,H). For both 50 and 100 SNPs windows, high frequency shared haplotypes were common for all six-row populations see for example Figure 5A, except for the Utah populations (Figures S10 and S11, Table S13 and S14). This contrasted with VT6 which showed very low degree of haplotype sharing, particularly with the various spring breeding populations (Figure 5B). Haplotype sharing for VT6 occurred primarily with the OR2 and OR6 populations; *i.e.*, with the only other programs with similar growth habit. However, OR2 and OR6 showed higher levels of allele frequency similarity with other breeding programs than VT6 (Figure S10J,K,N). These results are consistent with differentiation in allele frequency observed in *F*_ST_ comparisons between breeding populations (Table S15).

**Figure 5.**
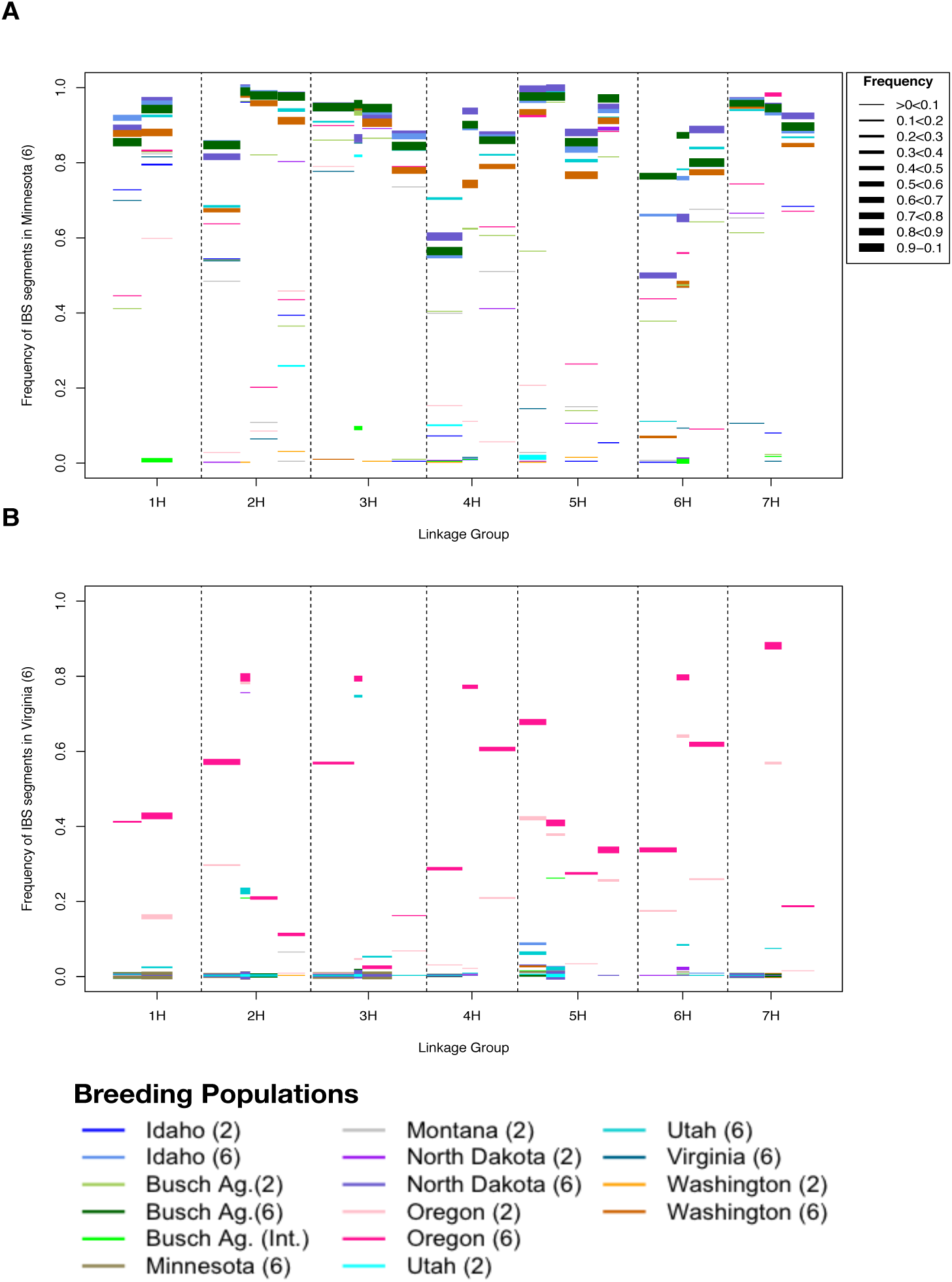
Frequency of identity by state haplotypes. **(A)** University of Minnesota breeding programs and **(B)** Virginia Polytechnic Institute and State University. Using 100 SNPs windows. Linkage groups are in the X-axis. The Y-axis is the frequency of the haplotype in the population indicated in the Y-axis label (“base population”). The color of the bars in the plot represent the distinct barley breeding programs that share the same segment with the “base population”. The width of each bar corresponds to the frequency of that haplotype in the compared populations.

For 50 SNPs windows, there were a high number of IBS segments shared between row types. However, when windows of 100 SNPs were considered, the majority of the IBS haplotypes were at low frequency (<20% frequency) in one of the two populations compared (Tables S13 and S14). An exception to this was the comparison between OR2 and OR6, where shared haplotypes were quite common. Sharing occurred at every 100 SNPs window, with shared haplotypes at frequencies as high as 90% in the two populations (Figure S11K) with an average frequency within populations of 0.46 and 0.58 (for OR2 and OR6 respectively) (Table S14). IBS with 50 SNPs windows for N2 and N6 span 94% of all windows, with shared haplotypes occurring at average frequencies of 22% and 50%, respectively (Figure S10,Table S13). Increasing the window size to 100 SNPs identified many fewer shared haplotypes between N2 and N6, leaving 70% of the genome shared at an average frequency of 8% in N2 and 35% in N6 (Figure S11, Table S14).

### Maximum likelihood tree of relatedness and migration

We determined that the tree topology that best describes the 16 populations separates the programs first by row-type followed by growth habit (Figure 6), consistent with the *F*_ST_ and PCA results, with the exception of UT2 and UT6 that are not separated by row-type. In the TreeMix analysis, the two Utah and winter populations were more similar to ancestral allele frequencies, while MN6 was the most diverged.

**Figure 6.**
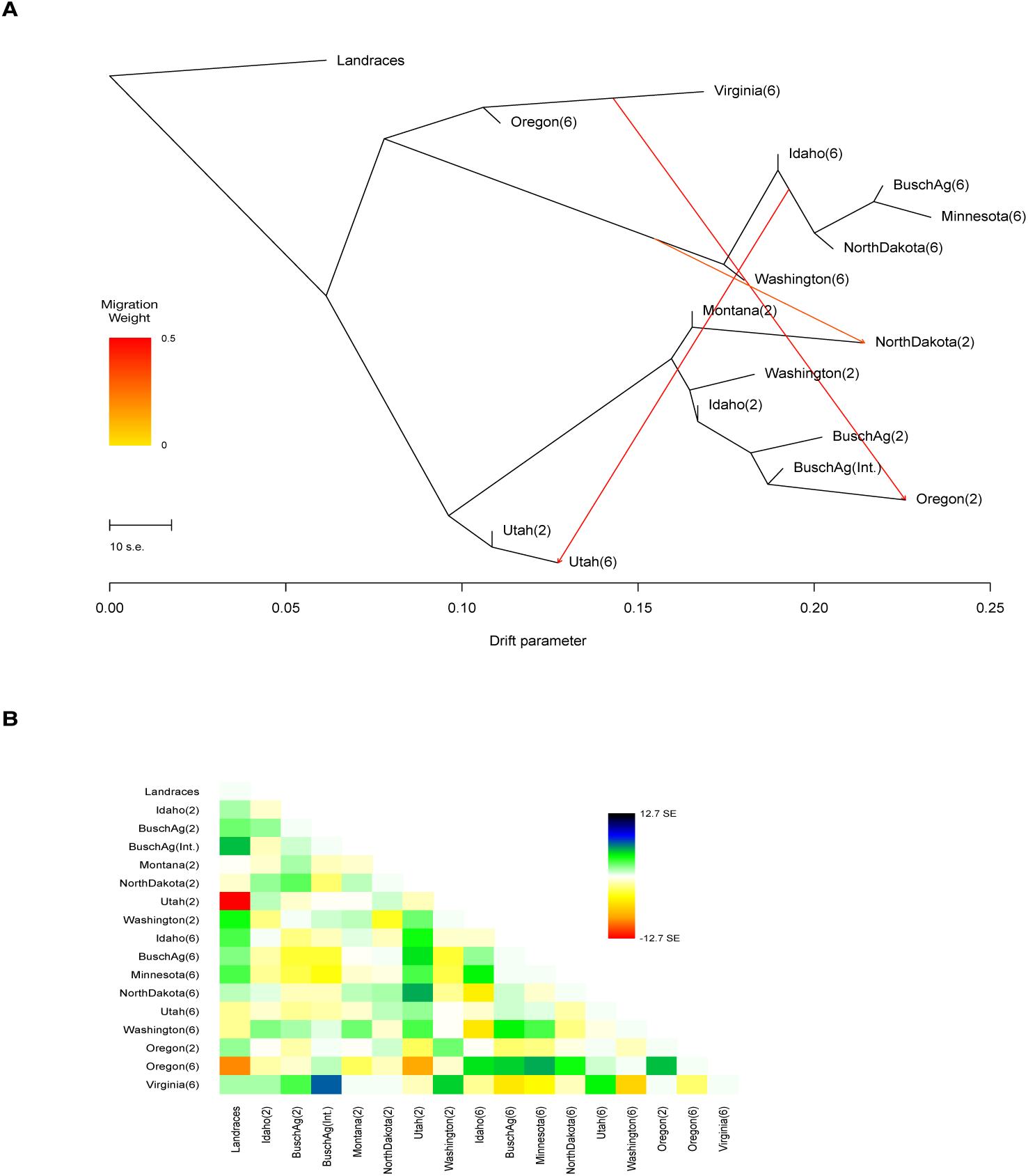
Tree of relatedness among North American breeding programs. **(A)** Plotted is the structure of the graph inferred by TreeMix for the 16 North American breeding populations, allowing three migration events. The arrows (migration events) are colored according to their weight. Horizontal branch length is proportional to the amount of genetic drift in the branch. The scale bar in the left shows ten times the average standard error of the entries in the sample covariance matrix. **(B)** Residual fit of inferred tree of relatedness. Residuals above zero represent populations that are more closely related to each other in the data than in the best-fit tree, thus are considered candidates for admixture events. Row-type for each population is presented in parenthesis.

Adding three migration events resulted in the lowest residuals and standard errors relative to trees with no migration, or one, two, four, or five migrations (Figure S12). With three migrations, we infer exchange between spring six-row programs and N2; VT6/OR6 and OR2, and spring six-row programs and UT6 (Figure 6).

Based on the *f*_4_-*test*, six different topologies yielded significant results suggesting gene flow between UT6 and spring six-row populations (Table S16), with z-scores significantly different from zero (significance at *p* <0.05). The *f*_*4*_-*test* for introgression between N2 and any spring six-row program, resulted in values significantly different from zero, consistent with gene flow. The *f*_*4*_-test did not support introgression between OR2 and VT6 and/or OR6 since none of the possible trees topologies involving these populations were significant.

## DISCUSSION

Comparative analyses of allele frequency differentiation and extend haplotype sharing in a sample of 3,613 barley accessions representing 16 barley breeding populations results in five primary conclusions: i) barley breeding populations in North America have strongly differentiated allele frequencies among breeding populations. Across programs, inflorescence type followed by growth habit account for the greatest proportion of variance in allele frequency, ii) a number of loci previously identified as major contributors to growth habit adaptation in barley are readily identifiable as outliers in allele frequency in our *F*_ST_ analyses, iii) we identify putative signals of recent and long-term selection similar in magnitude to previously isolated genes of known function, the average per SNP allele frequency divergence at the population level appears consistent with the dominant action of genetic drift and linked selection, and v) identification of populations are recently subject to exchange of genetic material. There are low levels of genetic exchange across inflorescence row-type boundaries.

### Identification of loci putatively subject to recent and long-term selection

Allele frequency differentiation sufficient to be detected in an *F*_ST_ outlier analysis is generally the result of long-term directional selection (Beaumont and Balding, 2004). The SNPs identified as outliers for spring versus winter growth habit include previously isolated genes of large effect, for example *Vrn-H1* and *Ppd-H1*, congruent with a previous *F*_ST_ analysis in European barleys (Comadran et al., 2012). Allele frequency differentiation between row-type, which explains a larger portion of the allele frequency divergence in our sample, identified two SNPs in the genomic region of genes contributing to row-type, however, SNPs within the well-characterized *Vrs1* locus were not identified as *F*_ST_ outliers (see below). Many additional *F*_ST_ outliers are newly identified as putative targets of selection.

Despite a relatively recent introduction to North America, multiple genes found to contribute to agronomic traits in Eurasia appear to recapitulate allele frequencies in current, highly structured, breeding populations. It bodes well for the potential to translate genetic results including efforts to identify particular causative genes and mutations across breeding populations and regions with distinct demographic and breeding histories when the phenotype is conferred by a single allele (as oppose to an allelic series, as for example, *Vrs1*).

PHS analysis in the North American breeding programs detected a series of loci that are putatively targets of long-term selection within barley breeding programs, comparable to findings in wheat (Cavanagh et al., 2013). The length of the haplotypes shared (average 81.38 cM) and their frequencies (average 25%) suggest that these events have taken place in recent generations so that recombination has not had time to break down the haplotypes. This analysis indicates that selection and linked selection are altering patterns of haplotype diversity across the genome. However, there are some chromosomes that have little or no evidence of selection in most recent generations (*e.g.*, linkage group 2H in BA2 and WA2. See Figure S8).

### Limitations of allele frequency comparisons

Despite having five SNPs positioned in the *Vrs1* gene (controlling the fertility of lateral spikelet) (Komatsuda and Mano, 2002), including one SNP (12_30900) variant co-segregating with the six-row phenotype, the comparison of allele frequencies differentiation between row-type fails to identify outliers in *Vrs1*. One important contributing factor may be that the six-row phenotype can result from multiple disruptions of the *Vrs1* gene (Komatsuda et al., 2007). The phenomenon of multiple mutational paths to the same phenotype has been identified as “functionally equivalent mutations” (Ralph and Coop, 2010) and can result in multiple variants at modest frequency controlling a phenotype. The maintenance of alleles of large effect at modest frequency in the population reduces the power to detect phenotypic associations, in part because any given variant explains a small portion of phenotypic variation (Thornton et al., 2013).

The BOPA SNP platform used here is largely derived from variants identified from cDNA libraries, from cultivated accessions (Close et al., 2009), and thus is highly enriched for SNPs in genic regions (Kono et al., 2013). However, SNPs genotyped within a locus can have limited correlation with a phenotype, depending on the haplotype on which they occur (see Nordborg and Tavaré, 2002for discussion of this issue). This may be the case for the previously cloned *Vrn-H3* where we see limited correlation between SNPs in this gene and allele frequency difference in the growth habit comparison. It should also be noted that the resequencing panel in which *Vrn-H3* alleles were found to be associated with the phenotype included only five spring and eight winter individuals (Yan et al., 2006). It is perhaps not surprising that sampling error (*i.e.,* small sample sizes) accounts for inconsistencies in the phenotype-genotype associations reported in previous studies and our study using a panel of >3,000 accessions.

### The effects of linked selection and drift in breeding programs

The derived site frequency spectrum in individual breeding population identifies an excess of rare and high frequency derived variants (Figure S5) with respect to neutral model of a population in equilibrium. During selection, alleles linked to the target variant can be carried to high frequencies. Therefore, an excess of high and rare frequency of derived variants can be associated with the effects of linked selection (Fay and Wu, 2000) as has been observed in a resequencing study of *Oryza sativa* (Caicedo et al., 2007). The PHS results presented here suggest considerable potential to detect the effect of linked selection, particularly when the selection was recent, impacting large genomic regions. One of the largest genomic region detected as an outlier in the PHS analysis is found in OR2, this haplotype involves 138.9 cM on linkage group 2H, observed at 11% frequency. Individuals carrying this haplotype derived from a two-row by six-row cross (Merlin x Strider) and were selected to recover the two-row phenotype (Dr. Hayes, personal communication). This resulted in a long shared haplotype centered on the *Vrs1* gene contributing to row-type (identified by SNPs 12_30901). Two of the SNPs that permitted the identification of the outlier haplotype occur within an iron homeostasis (*nashor 2*) and drought-stress tolerance genes (*TaGolS 2*).

Our comparisons of allele frequency differences among individual barley breeding populations (see Figure 4 and Table S15) suggests that genetic drift (and likely linked selection) play a major role in differentiation among populations. The cumulative effects of selection and genetic drift over several generations of breeding may result in reduced response to selection (Hanrahan et al., 1973), with larger effects of drift when the population is small and/or highly inbred (Robertson, 1960). The magnitude of drift suggests that the majority of barley breeding programs have small effective population sizes. Gerke *et al*. (2014) noted that genetic drift played a major role in changes in allele frequency over the history of maize breeding in North America. Although, in the larger partitions of barley populations, signals of directional selection are clearly evident at loci of large effect, drift may be extremely important to the loss of variation for variants of small effect (or for phenotypes determined by multiple genes), especially when the selective pressure is sporadic (*e.g*., drought tolerance).

### Gene flow among North American breeding programs

Gene flow between populations can be detected by the presence of large chromosomal regions in (admixture) linkage disequilibrium (Chakraborty and Smouse, 1988; Chakraborty and Weiss, 1988; Briscoe et al., 1994) or extended genomic regions of shared ancestry (Gusev et al., 2009; Gusev et al., 2012). Shorter shared haplotypes indicate shared ancestry a larger number of generation before present (more generations of recombination). Considering the high frequency of IBS segments (and associated low *F*_ST_) across the genome between populations of similar inflorescence type, we speculate that these segments could reflect a history of shared ancestry among these programs, possibly pre-dating the separation of breeding programs (Martin et al., 1991).

Gene flow that involves adaptive variation can result in differential retention of genomic segments in the recipient population, and can also be evident as shared genomic regions with reduced diversity (Hufford et al., 2013). For example, MN6 demonstrates a high degree of IBS with other six-row populations (Figure 6A), and these genomic regions have lower average pairwise diversity than other regions when the six-row populations are analyzed together (Figure S13).

With regard to IBS, VT6 is the most clearly differentiated from other North American populations, showing similarity only to the other winter programs, OR2 and OR6 (for 50 SNPs windows, Figure S10). Pairwise *F*_ST_ with other breeding programs and genetic differentiation as measured by PCA (Figure S2B) suggest that this isolation has been maintained over many generations. Large IBS segments (100 SNPs windows) shared between VT6 and OR2 and OR6 (Figure 5B) indicate that winter programs have recently exchanged genetic material. The demographic history inferred in the TreeMix analysis supports this relationship (Figure 6), particularly when invoking three migration events, the topology best supported by the data. However, the formal examination of migration using the *f*_*4*_-*test* failed to support gene flow among two-row and six-row winter programs, thus suggesting that haplotype similarities across row-types are due to ancestral history rather than recent introgression. In addition to the postulated shared ancestry, genetic similarity is expected between these programs under the premise that both Virginia and Oregon breeding programs have been subject to relative recent introgression from leaf rust resistance lines from the International Maize and Wheat Improvement Center (CIMMYT) (Dr. Griffey and Dr. Hayes, personal communication). Isolated populations, like VT6, can carry adaptive variants absent in other populations (Kristiansson et al., 2008). Therefore, VT6 could be used for the identification of new variants for yield or disease resistance, as well as a source of genetic variation to increase diversity in other breeding programs.

The high degree IBS segments shared between the Utah populations and most of the six-row populations yielded a significant signal of introgression based on the *f*_*4*_-*test*. It is possible that the signal detected here is due to the ongoing genetic exchange between UT2 and UT6 (as documented in pedigrees) since 2001 (Dr. David Hole, personal communication) and the genetic history shared between six-row barleys. This may also account for the elevated pairwise diversity in the Utah populations. Additionally, there is evidence of introgression between N2 and two six-row populations (WA6 and AB6), but not with N6. The introgression between the six-row populations and N2 are uncommon (Dr. Jerome Franckowiak, personal communication), thus this result needs more exploration.

Using the tree of relatedness as our hypothetical relationship, a necessary component of the *f*_*4*_-*test*, forces populations of the same row-type to be assumed as related by shared history. Therefore, gene flow within row-types is not investigated. Although, crosses across row-type are not common in barley breeding programs due to concerns of recombining desirable alleles with less favorable ones (Martin et al., 1991), we detect two instances of introgression across row-type that result in higher levels of genetic diversity as it was speculated by Martin *et al*. (Martin et al., 1991).

### Implications

Comparative population genetic approaches have the potential to uncover breeding history at a level of detail not previously possible. Our applications of allele frequency differentiation analyses in barley breeding programs are able to identify candidate genes or linked markers controlling major traits. Though the nature of the target of selection identified by individual SNPs is not always readily apparent, the identification of *F*_ST_ outliers reported here provides a reasonable first step toward the discovery of genes underlying agronomic adaptation or potential markers linked to these genes.

*F*_ST_ analyses also reveal the effect that linked selection and drift have in breeding programs. Taking into account that with any degree of linkage a proportion of the genetic variation may not be immediately available for selection (Hill and Robertson, 1966; Robertson, 1970); furthermore, beneficial variants could be in linkage with deleterious mutations (Felsenstein, 1974). Gains from selection in breeding programs will depend on the disassociation of these linkage blocks (Morrell et al., 2012; Rodgers-Melnick et al., 2015). This can be achieved by increasing the effective amount of recombination. Hill and Robertson (Hill and Robertson, 1966) suggested that a relaxed generation with a large number of contributing parents between each generation of selection could increase the amount of recombination needed to disassociate these variants.

### Caveats of the analysis

It is important to note that the results presented here are dependent on the samples submitted for genotyping (https://triticeaetoolbox.org) by each breeding program. Given that one of the goals of the Barley Coordinated Agricultural Project (http://www.barleycap.org) was to evaluate the diversity in the North American barley breeding program each program was encourage to submit representative lines. We assume that the data represent the diversity within each breeding programs and that the same sampling scheme across breeding programs is comparable. Deviations from these assumption could influence the results in four major ways: *i*) under estimation of the diversity within breeding programs; *ii*) over estimation of the role of drift in breeding populations due to reduced representation of the parental lines used in the breeding programs; *iii*) over or under estimation of allele frequency differentiation between partitions of the data; and *iv*) excess of shared haplotypes due to accessions highly related by pedigree (*e.g.,* sibs and half-sibs). With these caveats in mind, we made the best use of the dataset to uncover the underlying genetic response to breeder’s efforts. We encourage readers of this paper and barley breeders to evaluate the results with an understanding of the limitations of the sampling.

## ACKNOWLEDGEMENTS

The authors thank Michael Kantar, Ron Okagaki, and Paul Hoffman for helpful comments on an earlier version of the manuscript. We thank Yaniv Brandvain for helpul discussion about the manuscript. We thank Dr. David Hole, Dr. Patrick Hayes, and Dr. Jerome Franckowiak for sharing information about their breeding programs. This work was performed using computing resources at the University of Minnesota Supercomputing Institute. We acknowledge funding from the US Department of Agriculture National Institute for Food and Agriculture USDA NIFA 2011-68002-30029 (to P.L.M.).

## REFERENCES

Beaumont, M. A., and D. J. Balding, 2004 Identifying adaptive genetic divergence among populations from genome scans. Mol Ecol 13: 969–980.

Boyd, W. J. R., C. D. Li, C. R. Grime, M. Cakir, S. Potipibool, L. Kaveeta, S. Men, M. R. J. Kamali, A. R. Barr, and D. B. Moody, 2003 Conventional and molecular genetic analysis of factors contributing to variation in the timing of heading among spring barley (*Hordeum vulgare L.*) genotypes grown over a mild winter growing season. Crop Pasture Sci 54: 1277–1301.

Briscoe, D., J. C. Stephens, and S. J. O’Brien, 1994 Linkage disequilibrium in admixed populations: applications in gene mapping. J Hered 85: 59–63.

Caicedo, A. L., S. H. Williamson, R. D. Hernandez, A. Boyko, A. Fledel-Alon, T. L. York, N. R. Polato, K. M. Olsen, R. Nielsen, S. R. Mccouch, C. D. Bustamante, and M. D. Purugganan, 2007 Genomewide patterns of nucleotide polymorphism in domesticated rice. PLoS Genet 3: 1745–1756.

Cavalli-Sforza, L. L., 1966 Population structure and human evolution. Proc. R. Soc. Lond. B 362–379.

Cavanagh, C. R., S. Chao, S. Wang, B. E. Huang, S. Stephen, S. Kiani, K. Forrest, C. Saintenac, G. L. Brown-Guedira, and A. Akhunova, 2013 Genome-wide comparative diversity uncovers multiple targets of selection for improvement in hexaploid wheat landraces and cultivars. P Natl Acad Sci USA 110: 8057–8062.

Chakraborty, R., and P. E. Smouse, 1988 Recombination of haplotypes leads to biased estimates of admixture proportions in human populations. Proc Natl Acad Sci USA 85: 3071–3074.

Chakraborty, R., and K. M. Weiss, 1988 Admixture as a tool for finding linked genes and detecting that difference from allelic association between loci. Proc Natl Acad Sci U S A 85: 9119–9123.

Chang, C. C., C. C. Chow, L. C. Tellier, S. Vattikuti, S. M. Purcell, and J. J. Lee, 2015 Second-generation PLINK: rising to the challenge of larger and richer datasets. Gigascience 4: 7.

Close, T. J., P. R. Bhat, S. Lonardi, Y. Wu, N. Rostoks, L. Ramsay, A. Druka, N. Stein, J. T. Svensson, S. Wanamaker, S. Bozdag, M. L. Roose, M. J. Moscou, S. Chao, R. K. Varshney, P. Szucs, K. Sato, P. M. Hayes, D. E. Matthews, A. Kleinhofs, G. J. Muehlbauer, J. Deyoung, D. F. Marshall, K. Madishetty, R. D. Fenton, P. Condamine, A. Graner, and R. Waugh, 2009 Development and implementation of high-throughput SNP genotyping in barley. BMC Genomics 10: 582.

Comadran, J., J. R. Russell, A. Booth, A. Pswarayi, S. Ceccarelli, S. Grando, A. M. Stanca, N. Pecchioni, T. Akar, A. Al-Yassin, A. Benbelkacem, H. Ouabbou, J. Bort, F. A. Van Eeuwijk, W. T. Thomas, and I. Romagosa, 2011 Mixed model association scans of multienvironmental trial data reveal major loci controlling yield and yield related traits in *Hordeum vulgare* in Mediterranean environments. Theor Appl Genet 122: 1363–1373.

Comadran, J., B. Kilian, J. Russell, L. Ramsay, N. Stein, M. Ganal, P. Shaw, M. Bayer, W. Thomas, and D. Marshall, 2012 Natural variation in a homolog of *Antirrhinum* CENTRORADIALIS contributed to spring growth habit and environmental adaptation in cultivated barley. Nat Genet 44: 1388–1392.

Cuesta-Marcos, A., P. Szucs, T. J. Close, T. Filichkin, G. J. Muehlbauer, K. P. Smith, and P. M. Hayes, 2010 Genome-wide SNPs and re-sequencing of growth habit and inflorescence genes in barley: implications for association mapping in germplasm arrays varying in size and structure. BMC Genomics 11: ARTN 707.

Fang, Z., A. M. Gonzales, M. T. Clegg, K. P. Smith, G. J. Muehlbauer, B. J. Steffenson, and P. L. Morrell, 2014 Two genomic regions contribute disproportionately to geographic differentiation in wild barley. G3 4: 1193–1203.

Fang, Z., A. Eule-Nashoba, C. Powers, T. Y. Kono, S. Takuno, P. L. Morrell, and K. P. Smith, 2013 Comparative analyses identify the contributions of exotic donors to disease resistance in a barley experimental population. G3 3: 1945–1953.

Fay, J. C., and C. I. Wu, 2000 Hitchhiking under positive Darwinian selection. Genetics 155: 1405–1413.

Felsenstein, J., 1974 The evolutionary advantage of recombination. Genetics 78: 737–756.

Gerke, J., J. Edwards, G. Ke, J. Ross-Ibarra, and M. D. Mcmullen, 2014 The genomic impacts of drift and selection for hybrid performance in maize. arXiv:1307.7313

Goudet, J., 2005 Hierfstat, a package for R to compute and test hierarchical F statistics. Mol Ecol Notes 5: 184–186.

Gusev, A., P. F. Palamara, G. Aponte, Z. Zhuang, A. Darvasi, P. Gregersen, and I. Pe’Er, 2012 The architecture of long-range haplotypes shared within and across populations. Mol Biol Evol 29: 473–486.

Gusev, A., J. K. Lowe, M. Stoffel, M. J. Daly, D. Altshuler, J. L. Breslow, J. M. Friedman, and I. Pe’Er, 2009 Whole population, genomewide mapping of hidden relatedness. Genome Res 19: 318–326.

Haake, V., D. Cook, J. L. Riechmann, O. Pineda, M. F. Thomashow, and J. Z. Zhang, 2002 Transcription factor CBF4 is a regulator of drought adaptation in Arabidopsis. Plant Physiol 130: 639–648.

Hamblin, M. T., T. J. Close, P. R. Bhat, S. Chao, J. G. Kling, K. J. Abraham, T. Blake, W. S. Brooks, B. Cooper, and C. A. Griffey, 2010 Population structure and linkage disequilibrium in US barley germplasm: implications for association mapping. Crop Sci 50: 556–566.

Hanrahan, J. P., E. J. Eisen, and J. E. Lagates, 1973 Effects of population size and selection intensity of short-term response to selection for postweaning gain in mice. Genetics 73: 513–530.

Herbik, A., G. Koch, H. P. Mock, D. Dushkov, A. Czihal, J. Thielmann, U. W. Stephan, and H. BäUmlein, 1999 Isolation, characterization and cDNA cloning of nicotianamine synthase from barley. A key enzyme for iron homeostasis in plants. Eur J Biochem 265: 231–239.

Hill, W. G., and A. Robertson, 1966 The effect of linkage on limits to artificial selection. Genet Res 8: 269–294.

Horton, M. W., A. M. Hancock, Y. S. Huang, C. Toomajian, S. Atwell, A. Auton, N. W. Muliyati, A. Platt, F. G. Sperone, B. J. VilhjÁLmsson, M. Nordborg, J. O. Borevitz, and J. Bergelson, 2012 Genome-wide patterns of genetic variation in worldwide Arabidopsis thaliana accessions from the RegMap panel. Nat Genet 44: 212–216.

Hudson, R. R., K. Bailey, D. Skarecky, J. Kwiatowski, and F. J. Ayala, 1994 Evidence for positive selection in the superoxide dismutase (Sod) region of *Drosophila melanogaster*. Genetics 136: 1329–1340.

Hufford, M. B., P. Lubinksy, T. Pyhajarvi, M. T. Devengenzo, N. C. Ellstrand, and J. Ross-Ibarra, 2013 The genomic signature of crop-wild introgression in maize. PLoS Genet 9: e1003477.

Innan, H., K. Zhang, P. Marjoram, S. TavarÉ, and N. A. Rosenberg, 2005 Statistical tests of the coalescent model based on the haplotype frequency distribution and the number of segregating sites. Genetics 169: 1763–1777.

Jones, H., F. J. Leigh, I. Mackay, M. A. Bower, L. M. J. Smith, M. P. Charles, G. Jones, M. K. Jones, T. A. Brown, and W. Powell, 2008 Population-based resequencing reveals that the flowering time adaptation of cultivated barley originated east of the Fertile Crescent. Mol Biol Evol 25: 2211–2219.

Karsai, I., P. Szucs, B. Koszegi, P. M. Hayes, A. Casas, Z. Bedo, and O. Veisz, 2008 Effects of photo and thermo cycles on flowering time in barley: agenetical phenomics approach. J Exp Bot 59: 2707–2715.

Kearse, M., R. Moir, A. Wilson, S. Stones-Havas, M. Cheung, S. Sturrock, S. Buxton, A. Cooper, S. Markowitz, C. Duran, T. Thierer, B. Ashton, P. Meintjes, and A. Drummond, 2012 Geneious Basic: an integrated and extendable desktop software platform for the organization and analysis of sequence data. Bioinformatics 28: 1647–1649.

Keinan, A., J. C. Mullikin, N. Patterson, and D. Reich, 2007 Measurement of the human allele frequency spectrum demonstrates greater genetic drift in East Asians than in Europeans. Nat Genet 39: 1251–1255.

Kimura, M., 1983 The Neutral Theory of Molecular Evolution. University Press, Cambridge, United Kingdom, Cambridge.

Komatsuda, T., and Y. Mano, 2002 Molecular mapping of the intermedium spike-c (int-c) and non-brittle rachis 1 (btr1) loci in barley (*Hordeum vulgare L.*). Theor Appl Genet 105: 85–90.

Komatsuda, T., M. Pourkheirandish, C. He, P. Azhaguvel, H. Kanamori, D. Perovic, N. Stein, A. Graner, T. Wicker, and A. Tagiri, 2007 Six-rowed barley originated from a mutation in a homeodomain-leucine zipper I-class homeobox gene. Proc Natl Acad Sci USA 104: 1424–1429.

Kono, T. J. Y., K. Seth, and J. A. Poland, 2013 SNPMeta: SNP annotation and SNP Metadata collection without a reference genome. Mol Ecol

Kristiansson, K., J. Naukkarinen, and L. Peltonen, 2008 Isolated populations and complex disease gene identification. Genome Biol 9: 109.

Laurie, D. A., N. Pratchett, J. W. Snape, and J. H. Bezant, 1995 RFLP mapping of five major genes and eight quantitative trait loci controlling flowering time in a winter× spring barley (*Hordeum vulgare L*.) cross. Genome 38: 575–585.

Lewontin, R. C., and J. Krakauer, 1973 Distribution of gene frequency as a test of the theory of the selective neutrality of polymorphisms. Genetics 74: 175–195.

Lister, D. L., S. Thaw, M. A. Bower, H. Jones, M. P. Charles, G. Jones, L. M. J. Smith, C. J. Howe, T. A. Brown, and M. K. Jones, 2009 Latitudinal variation in a photoperiod response gene in European barley: insight into the dynamics of agricultural spread from ‘historic’ specimens. J Archaeol Sci 36: 1092–1098.

Marchini, J., 2013 Statistical and Population Genetics.

Martin, J. M., T. K. Blake, and E. A. Hockett, 1991 Diversity among North American spring barley cultivars based on coefficients of parentage. Crop Sci 31:1131–1137.

Mayer, K. F., R. Waugh, P. Langridge, T. J. Close, R. P. Wise, A. Graner,T. Matsumoto, K. Sato, A. Schulman, G. J. Muehlbauer, et. al., 2012 A physical, genetic and functional sequence assembly of the barley genome. Nature 491: 711–716.

Morrell, P. L., E. S. Buckler, and J. Ross-Ibarra, 2012 Crop genomics: advances and applications. Nat Rev Genet 13: 85–96.

Muñoz-Amatriaín, M., A. Cuesta-Marcos, J. B. Endelman, J. Comadran, J. M. Bonman, H. E. Bockelman, S. Chao, J. Russell, R. Waugh, and P. M. Hayes, 2014 The USDA barley core collection: Genetic diversity, population structure, and potential for genome-wide association studies. PLoS One 9: e94688.

Muñoz-Amatriaín, M., M. J. Moscou, P. R. Bhat, J. T. Svensson, J. Bartoš, P. Suchánková, H. Šimková, T. R. Endo, R. D. Fenton, and S. Lonardi, 2011 An improved consensus linkage map of barley based on flowsortedchromosomes and single nucleotide polymorphism markers. Plant Genome 4: 238–249.

Nicholson, G., A. V. Smith, F. Jónsson, Ó. Gústafsson, K. Stefánsson, and P. Donnelly, 2002 Assessing population differentiation and isolation from single-nucleotide polymorphism data. J R Stat Soc Series B Stat Methodol 64: 695–715.

Nielsen, R., I. Hellmann, M. Hubisz, C. Bustamante, and A. G. Clark, 2007 Recent and ongoing selection in the human genome. Nat Rev Genet 8: 857–868.

Nordborg, M., and S. Tavaré, 2002 Linkage disequilibrium: what history has to tell us. Trends Genet 18: 83–90.

Patterson, N., A. L. Price, and D. Reich, 2006 Population structure and eigenanalysis. PLoS Genet 2: e190.

Pickrell, J. K., and J. K. Pritchard, 2012 Inference of population splits and mixtures from genome-wide allele frequency data. PLoS Genet 8: e1002967.

Poets, A. M., Z. Fang, M. T. Clegg, and Morrell, 2015 Barley landraces are characterized by geographically heterogeneous genomic origins. Genome Biol 16: 173.

Pritchard, J. K., J. K. Pickrell, and G. Coop, 2010 The genetics of human adaptation: hard sweeps, soft sweeps, and polygenic adaptation. Curr Biol 20: R208–15.

Team, R. D. C., 2012 R: A language and environment for statistical computing. R Foundation for Statistical Computing, Vienna, Austria.

Ralph, P., and G. Coop, 2010 Parallel adaptation: one or many waves of advance of an advantageous allele? Genetics 186: 647–668.

Ramsay, L., J. Comadran, A. Druka, D. F. Marshall, W. T. Thomas, M. Macaulay, K. Mackenzie, C. Simpson, J. Fuller, N. Bonar, P. M. Hayes, U. Lundqvist, J. D. Franckowiak, T. J. Close, G. J. Muehlbauer, and R. Waugh, 2011 INTERMEDIUM-C, a modifier of lateral spikelet fertility in barley, is an ortholog of the maize domestication gene TEOSINTE BRANCHED 1. Nat Genet 43: 169–172.

Reich, D., K. Thangaraj, N. Patterson, A. L. Price, and L. Singh, 2009 Reconstructing Indian population history. Nature 461: 489–494.

Robertson, A., 1970 A theory of limits in artificial selection with many linked loci, pp. 246–288 in Mathematical topics in population genetics, edited by Springer,

Robertson, A., 1960 A theory of limits in artificial selection. Proc R Soc B 153: 234–249.

Rodgers-Melnick, E., P. J. Bradbury, R. J. Elshire, J. C. Glaubitz, C. B. Acharya, S. E. Mitchell, C. Li, Y. Li, and E. S. Buckler, 2015 Recombination in diverse maize is stable, predictable, and associated with genetic load. Proc Natl Acad Sci U S A 112: 3823–3828.

Ross-Ibarra, J., P. L. Morrell, and B. S. Gaut, 2007 Plant domestication, a unique opportunity to identify the genetic basis of adaptation. P Natl Acad Sci USA 104: 8641–8648.

Scheet, P., and M. Stephens, 2006 A fast and flexible statistical model for largescale population genotype data: applications to inferring missing genotypes and haplotypic phase. Am J Hum Gen 78: 629–644.

Skinner, J. S., J. Von Zitzewitz, P. Szucs, L. Marquez-Cedillo, T. Filichkin, K. Amundsen, E. J. Stockinger, M. F. Thomashow, T. H. Chen, and P. M. Hayes, 2005 Structural, functional, and phylogenetic characterization of a large CBF gene family in barley. Plant Mol Biol 59: 533–551.

Slatkin, M., 1987 Gene flow and the geographic structure of natural populations. Science 236: 787–792.

Taji, T., C. Ohsumi, S. Iuchi, M. Seki, M. Kasuga, M. Kobayashi, K. Yamaguchi-Shinozaki, and K. Shinozaki, 2002 Important roles of drought- and cold-inducible genes for galactinol synthase in stress tolerance in Arabidopsis thaliana. Plant J 29: 417–426.

Thornton, K., 2003 libsequence: a C++ class library for evolutionary genetic analysis. Bioinformatics 19: 2325–2327.

Thornton, K. R., A. J. Foran, and A. D. Long, 2013 Properties and modeling of GWAS when complex disease risk is due to non-complementing, deleterious mutations in genes of large effect. PLoS Genet 9: e1003258.

Toomajian, C., T. T. Hu, M. J. Aranzana, C. Lister, C. Tang, H. Zheng, K. Zhao, P. Calabrese, C. Dean, and M. Nordborg, 2006 A nonparametric test reveals selection for rapid flowering in the *Arabidopsis* genome. PLoS Biol 4: e137.

Turck, F., F. Fornara, and G. Coupland, 2008 Regulation and identity of florigen: FLOWERING LOCUS T moves center stage. Annu. Rev. Plant Biol. 59: 573–594.

Turner, A., J. Beales, S. Faure, R. P. Dunford, and D. A. Laurie, 2005 The pseudo-response regulator Ppd-H1 provides adaptation to photoperiod in barley. Science 310: 1031–1034.

Von Zitzewitz, J., P. Szucs, J. Dubcovsky, L. Yan, E. Francia, N. Pecchioni, A. Casas, T. H. Chen, P. M. Hayes, and J. S. Skinner, 2005 Molecular and structural characterization of barley vernalization genes. Plant Mol Biol 59: 449–467.

Wang, H., K. P. Smith, E. Combs, T. Blake, R. D. Horsley, and G. J. Muehlbauer, 2012 Effect of population size and unbalanced data sets on QTL detection using genome-wide association mapping in barley breeding germplasm. Theor Appl Genet 124: 111–124.

Weaver, J. C., 1943 Barley in the United States: A Historical Sketch. Geographical Review 33: 56–73.

Weaver, J. C., 1944 United States Malting Barley Production. Taylor & Francis, Ltd. on behalf of the Association of American Geographers.

Weaver, J. C., 1950 American barley production: A study in agricultural Geography. Burgess Pub. Co., Minneapolis, MN.

Weir, B. S., and C. C. Cockerham, 1984 Estimating F-statistics for the analysis of population structure. Evolution 38: 1358–1370.

Wiebe, G. A., and D. A. Reid, 1961 Classification of barley varieties growing in the United States and Canada in 1958. U.S. Department of Agriculture 210.

Yan, L., D. Fu, C. Li, A. Blechl, G. Tranquilli, M. Bonafede, A. Sanchez, M. Valarik, S. Yasuda, and J. Dubcovsky, 2006 The wheat and barley vernalization gene VRN3 is an orthologue of FT. Proc Natl Acad Sci U S A 103: 19581–19586.

Youssef, H. M., R. Koppolu, and T. Schnurbusch, 2012 Re-sequencing of vrs1 and int-c loci shows that labile barleys (*Hordeum vulgare convar. labile*) have a six-rowed genetic background. Genet Resour Crop Ev 59: 1319–1328.

Zhou, H., G. Muehlbauer, and B. N. Steffenson, 2012 Population structure and linkage disequilibrium in elite barley breeding germplasm from the United States. J Zhejiang Univ Sci B 13: 438–451.

